# SARS-CoV-2 variants show a gradual declining pathogenicity and pro-inflammatory cytokine spur, an increasing antigenic and antiinflammatory cytokine induction, and rising structural protein instability

**DOI:** 10.1101/2022.02.15.480592

**Authors:** Debmalya Barh, Sandeep Tiwari, Lucas Gabriel Rodrigues Gomes, Cecília Horta Ramalho Pinto, Bruno Silva Andrade, Shaban Ahmad, Alaa A. A. Aljabali, Khalid J. Alzahrani, Hamsa Jameel Banjer, Sk. Sarif Hassan, Murtaza Tambuwala, Elrashdy M. Redwan, Khalid Raza, Vasco Azevedo, Kenneth Lundstrom, Vladimir N. Uversky

## Abstract

Hyper-transmissibility with decreased disease severity are typical characteristics of Omicron variant. To understand this phenomenon, we used various bioinformatics approaches to analyze randomly selected genome sequences (one each) of the Gamma, Delta, and Omicron variants submitted to NCBI from 15 to 31 December 2021. We show that: (i) Pathogenicity of SARS-CoV-2 variants decreases in the order: Wuhan > Gamma > Delta > Omicron; however, the antigenic property follows the order: Omicron > Gamma > Wuhan > Delta. (ii) Omicron Spike RBD has lower pathogenicity but higher antigenicity than other variants. (iii) Decreased disease severity by Omicron variant may be due to its decreased pro-inflammatory and IL-6 stimulation and increased IFN-γ and IL-4 induction efficacy. (iv) Mutations in N protein are associated with decreased IL-6 induction and human DDX21-mediated increased IL-4 production in Omicron. (v) Due to mutations, the stability of S, M, N, and E proteins decreases in the order: Omicron > Gamma > Delta > Wuhan. (vi) Stronger Spike RBD-*h*ACE2 binding in Omicron is associated with increased transmissibility. However, the lowest stability of the Omicron Spike protein makes Spike RBD-*h*ACE2 interaction weak for systemic infection and for causing severe disease. Finally (vii), the highest instability of Omicron E protein may also be associated with decreased viral maturation and low viral load leading to less severe disease and faster recovery. Our method may be used for other similar viruses, and these findings will contribute to the understanding of the dynamics of SARS-CoV-2 variants and the management of emerging variants.

## INTRODUCTION

According to a recent report, in the USA, the cases of the Omicron (B.1.1.529) variant of SARS-CoV-2 have risen significantly compared to the Delta (B.1.617.2) or pre-Delta variants. The infection rate is particularly high in the age group of 18-49 years, and Omicron is highly transmissible even among fully vaccinated adults, where it is 2.7 to 3.7 times as infectious as the Delta variant. However, the death rate is low in the case of Omicron compared to the other variants [1]. Similar findings of high transmission rate and decreased disease severity were also reported in other countries, including South Africa, where the Omicron variant was first reported [2,3]. The reported death rate in South Africa from Wuhan, Delta, and Omicron are 19.7%, 29.1%, and 2.7%, respectively [4]. A similar low death rate was also seen in the USA and other countries [1]. However, the cause of high transmission rate and decreased disease severity of the Omicron variant is still fully not understood.

Reports suggest that Omicron can escape from antibody neutralization and vaccine protection due to mutations in its Spike (S) protein responsible for high transmission, and the attenuated replication of Omicron is associated with the decreased disease severity and death [5,6]. In most cases, the Spike (S) receptor-binding domain (RBD) and Angiotensin-converting enzyme 2 (*h*ACE2) interaction is in the focus of studies targeted at understanding the increased transmissibility of the Omicron variant, where some researches have shown that the Omicron RBD strongly binds to *h*ACE2 [7–9]. However, some other reports suggest a low affinity of the Omicron RBD to *h*ACE2 [10,11]. Therefore, further research is needed to understand the strength of the Omicron RBD - *h*ACE2 interaction and the stability of this complex.

To understand the features of Omicron that allow it to infect a wide range of age groups, we reviewed the literature to analyze why the other pre-Omicron variants of SARS-CoV-2 are not infecting or causing severe disease in young adults. From the immunological point of view, there are many possibilities why COVID-19 is less severe in young adults [12]. Chitinase 3-like-1 protein (CHI3L1) stimulates the expression of *h*ACE2 and viral Spike protein priming proteases (SPP) in older adults and comorbid conditions. This increased expression and availability of *h*ACE2 correlate with increased transmission of SARS-CoV-2 and severity from COVID-19 [13]. Another report indicates that high serum concentrations of interleukin-17A (IL-17A) and type II interferon (IFN-γ) in children and young adults provide an innate immunity against different SARS-CoV-2 strains [14,15]. It is also reported that children under 15 years of age show increased expression of type III interferon (IFN-λ1) in nasopharyngeal mucosa upon SARS-CoV-2 infection that prevents virus entry into the body [16].

On the other hand, the S and Nucleocapsid (N) proteins of SARS-CoV-2 induce antiinflammatory IFN-γ production in the host upon infection [17,18]. Further, the nonstructural protein-1 (NSP1) and N protein block the type I interferon (IFN-β) induction in the host and attenuate antiviral immune responses [19]. SARS-CoV-2 ORF8 activates the IL-17 signaling pathway and induces the secretion of inflammatory factors [20,21]. ORF8 also down-regulates MHC-I through lysosomal degradation and is involved in the SARS-CoV-2 immune evasion [22]. Furthermore, ORF8 attenuates the IFN-γ mediated antiviral responses in COVID-19 [23]. The membrane glycoprotein (M) of SARS-CoV-2 suppresses the expression and activity of IFN-λ1 through inhibition of the RIG-I/MDA-5 pathway [24]. Therefore, a wide range of hostpathogen protein-protein interactions modulates the host immune response in SARS-CoV-2 infection and disease severity.

Since the SARS-CoV-2 is a rapidly mutating virus with varying degrees of transmission and disease severity abilities, the mutations in the Omicron variant are responsible for its high transmission and reduced disease severity. In this context, we aimed to analyze the structural proteins (S, M, N, and E) and pathogenicity-associated important accessory proteins (ORF3a, ORF7a, ORF7b, ORF8, and ORF10) [21,25] of Omicron for a better understanding of the role these proteins in providing higher transmission and decreased disease severity/death of Omicron as compared to other SARS-CoV-2 variants of concern.

## METHODS

### Variants, genomes, and mutation mapping

We considered four SARS-CoV-2 variants in our analysis. Wuhan (wild-type SARS-CoV-2) (RefSeq: NC_045512.2) and three variants, whose sequences were submitted to the NCBI GenBank between December 15 and 31, 2021 and were randomly selected. The variants and their genomes are: Gamma (P.1) (B.1.1.28.1) accession no: MZ477769.1 (submission date: 31 December, 2021), Delta (B.1.617.2) accession no: OL966477.1 (submission date: 22 December, 2021), and Omicron (B.1.1.529) accession no: OL901854.1 (submission date: 17 December, 2021). The genome and proteome sequences were checked for their completeness. The structural and non-structural protein sequences were identified from these genomes for each variant. Multiple sequence alignment (MSA) was performed using Clustal Omega [26], and the mutations were identified based on the RefSeq Wuhan (wild-type) sequence. Jalview V.2 [27] was used to visualize the MSA results.

### Prediction of pathogenic and antigenic properties

We utilized the MP3 tool [28] with default parameters to predict the pathogenicity scores using the genome sequence of each of the selected variants. Hybrid results (SVM+HMM) were considered, and the SVM scores were used in the analysis. We individually calculated the pathogenic score for each protein of interest: ORFab, full-length S protein, Spike-RBD (amino acid residues 331 to 524 of S protein) [29–31], M, N, E, ORF3a, ORF7a, ORF7b, ORF8, and ORF10. The cumulative and average scores of all these 11 proteins for each variant were also calculated. For antigenic property analysis, amino acid sequences of each of these proteins from each variant were used, and the VaxiJen v2.0 server [32] with the threshold of 0.4 was applied. The same calculation made for pathogenicity prediction was also applied for antigenic property assessment.

### Prediction of cytokine and interleukin producing peptides

We predicted pro-inflammatory inducing peptides using the PIP-EL web server with its default parameters [33]. IFN-γ producing peptides were predicted using the IFNepitope server [34], selecting scan, motif, SVM hybrid method, and IFN-γ versus non-IFN-γ option with other default parameters. IL4pred [35] was used to predict IL-4 inducing peptides. We selected protein mapping and Hybrid (SVM + Merci motif) based prediction methods and kept other parameters as default. For prediction of IL-6 inducing peptides, we considered IL-6Pred [36], selecting protein scan and Random Forest (RF)-based prediction methods keeping other parameters as default. Finally, we used IL17eScan [37] to identify the IL-17 inducing peptides selecting protein scan module and DPC-based model. For selecting the final epitope-related calculations, we used threshold value (score) >0.05 for IFN-γ, IL-4, and IL-17 inducing peptides, and for IL-6 producing, we used the cut-off value as >0.3. The number of positive epitopes generated and the cumulative or average SVM score of the total number of positive epitopes at the set cut off for full-length S, Spike-RBD, M, N, E, ORF3a, ORF7a, ORF7b, ORF8, and ORF10 proteins were used for further calculations. The total number of positive epitopes and their cumulative or average SVM scores were used for the final results for each variant.

### Identification of immune-associated human proteins that interact with ORF8, M, and N of SARS-CoV-2

The interacting human protein partners of ORF8, M, and N proteins of SARS-CoV-2 as described by Gordon *et al*. [38] and Enrichr-based Gene set enrichment analysis (GSEA) [39] were used. To investigate the immune-modulating pathway for each of these SARS-CoV-2 proteins, we used the corresponding interacting human proteins in Enrichr. Under disease/drugs, LINCS L1000 ligand perturbations database [40]-based results were used to provide cytokine, interleukin, and interferon related regulation information. Thirty, 15, and 47 human proteins that interact with M, N, and ORF8 of SARS-CoV-2, respectively, according to Gordon *et al.* [38] were used in this analysis. We identified the key integrating human proteins modulating cytokine, interleukin, and interferon-related pathways upon interacting with ORF8, M, and N of SARS-CoV-2. Further, based on protein-protein docking using the HDOCK server [41], we predicted the mutational effects of ORF8, M, and N on binding activity and their possible input on regulating these cytokines interleukin and interferon-related pathways for each SARS-CoV-2 variant.

### Protein 3D structures

Crystal structures of Spike-*h*ACE2 complexes for Wuhan (PDB: 6M0J), Gamma (P.1) (PDB: 7EKC), Delta (PDB: 7V8B), and Omicron (PDB: 7T9L) variants were retrieved from the PDB database (https://www.rcsb.org). Other crystal structures used are human DExD-box helicase 21 (DDX21) (PDB: 6L5N), human G3BP stress granule assembly factor 1 (G3BP1) (PDB: 4FCJ), human Gamma-Glutamyl Hydrolase (GGH) (PDB: 1L9X), and SARS-CoV-2 ORF8 protein (PDB: 7F5F). For human Stomatin (hSTOM) (UniProtKB: P27105), AlphaFold based structure AF-P27105-F1 available in UniProt was used. SARS-CoV-2 M and N protein sequences were identified from the specific genomes we have used. The 3D models of M and N were obtained by using the RaptorX server [42], followed by the GalaxyRefine server [43]. We also used the AlphaFold [44] and SWISS-MODEL servers [45] to model the mutant proteins. The SAVES v6.0 server (https://saves.mbi.ucla.edu) based PROCHECK tool [46] was used to analyze the stereochemical quality of the modelled protein structures, and the best models were selected based on the Ramachandran plot parameters (> 90% residues in the most favored region).

### Protein-protein docking for binding strength calculation

The HDOCK server [41] was used for protein-protein docking. The UCSF Chimera X program [47] was used for 3D structure analysis and protein-protein 2D interaction maps were generated using LigPlot+ v.2.2 [48]. We also used the commercial version of Schrödinger Platform (https://www.schrodinger.com) for a second line validation. For Spike-*h*ACE2 analysis, for each variant, the crystal structure was cleaned for water molecules, ligands, ions, and crystallographic artifacts using Chimera X, and then the interactions / interacting residues between the Spike-RBD and *h*ACE2 were mapped using LigPlot+ v.2.2/ Schrödinger Platform. Next, we applied the HDOCK server and Schrödinger Platform and used the identified residues for protein-protein docking. The resultant best-docked complex was selected based on docking score and and H-bonds to understand the binding strength.

We also used HDOCK server and Schrödinger Platform to understand the interactions between (i) human STOM and four SARS-C0V-2 M protein variants, (ii) human G3BP1 and four SARS-CoV-2 N protein variants, (iii) human DDX21 and four SARS-CoV-2 N protein variants, and (iv)human STOM and four SARS-CoV-2 ORF8 protein variants. We performed blind docking for these interactions. Best docked complex was selected based on docking score and H-bonds, and LigPlot+ v.2.2 or Schrödinger Platform was used to identify the binding residues.

### Analysis of the effect of mutations on protein-protein interaction

Folding free energy (ΔΔG) -based possible effect of SARS-CoV-2 mutations in corresponding protein stability was performed by DynaMut2 [49]. The 3D structures of the Wuhan (wild-type) variant of SARS-CoV-2 proteins were used to predict the stability of the mutant proteins of other variants.

## RESULTS

### Omicron shows decreased pathogenicity and increased antigenic potential

Our analysis suggested a gradual decrease in pathogenicity and increased antigenicity as the new SARS-CoV-2 variants evolved. The overall pathogenic potential of Omicron is nearly 54% lower than that of the Wuhan variant, approximately 50% lower than the Gamma (P.1), and 20 % of the Delta strain (**Fig 1 A, C**). In the context of only the structural proteins, we observed the same trend; i.e., the order of pathogenicity is Wuhan > Gamma > Delta > Omicron (**Fig 1B**). The structural protein-based pathogenicity of Omicron is approximately 38% lower than that of Wuhan, 37% lower than Gamma, and 32% lower than the Delta variant (**Fig 1B, C**). It is important to note that, although the pathogenicity of spike RBD in Omicron is showing a negative result, it has very low pathogenicity. On the other hand, although the Omicron E protein shows a positive pathogenic value, it is less pathogenic compared to the E protein from other strains as per the MP3 hybrid prediction (**Fig 1A**). However, these values do not affect the overall low pathogenicity of the Omicron strain (**Fig 2A**).

**Fig 1:**
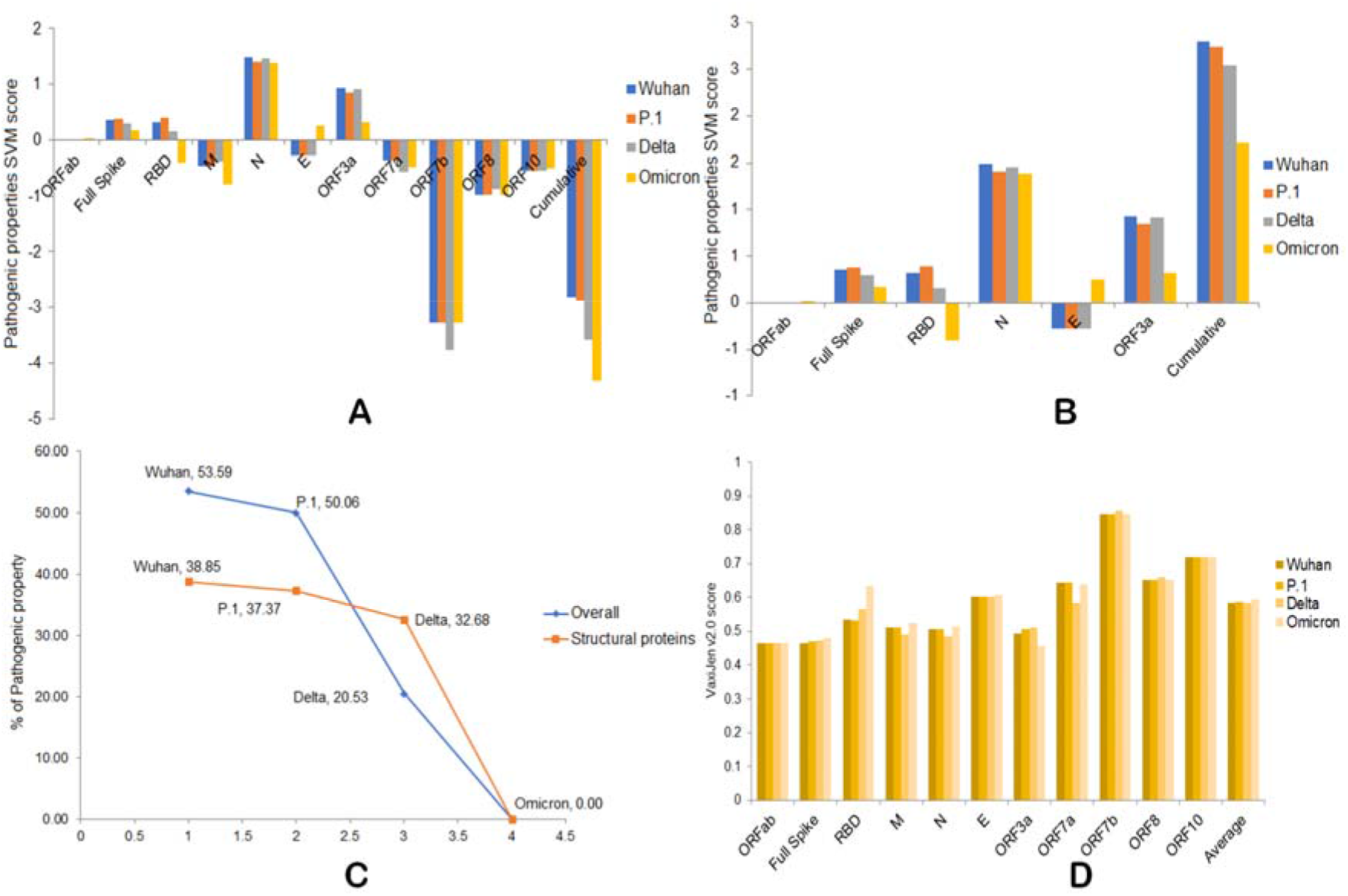
A) Overall pathogenic properties of the four variants. B) Overall pathogenic properties of structural proteins from the four variants. C) The percentage of decreased pathogenicity of the four variants as compared to Omicron. D) Antigenic properties of four variants.

**Fig 2:**
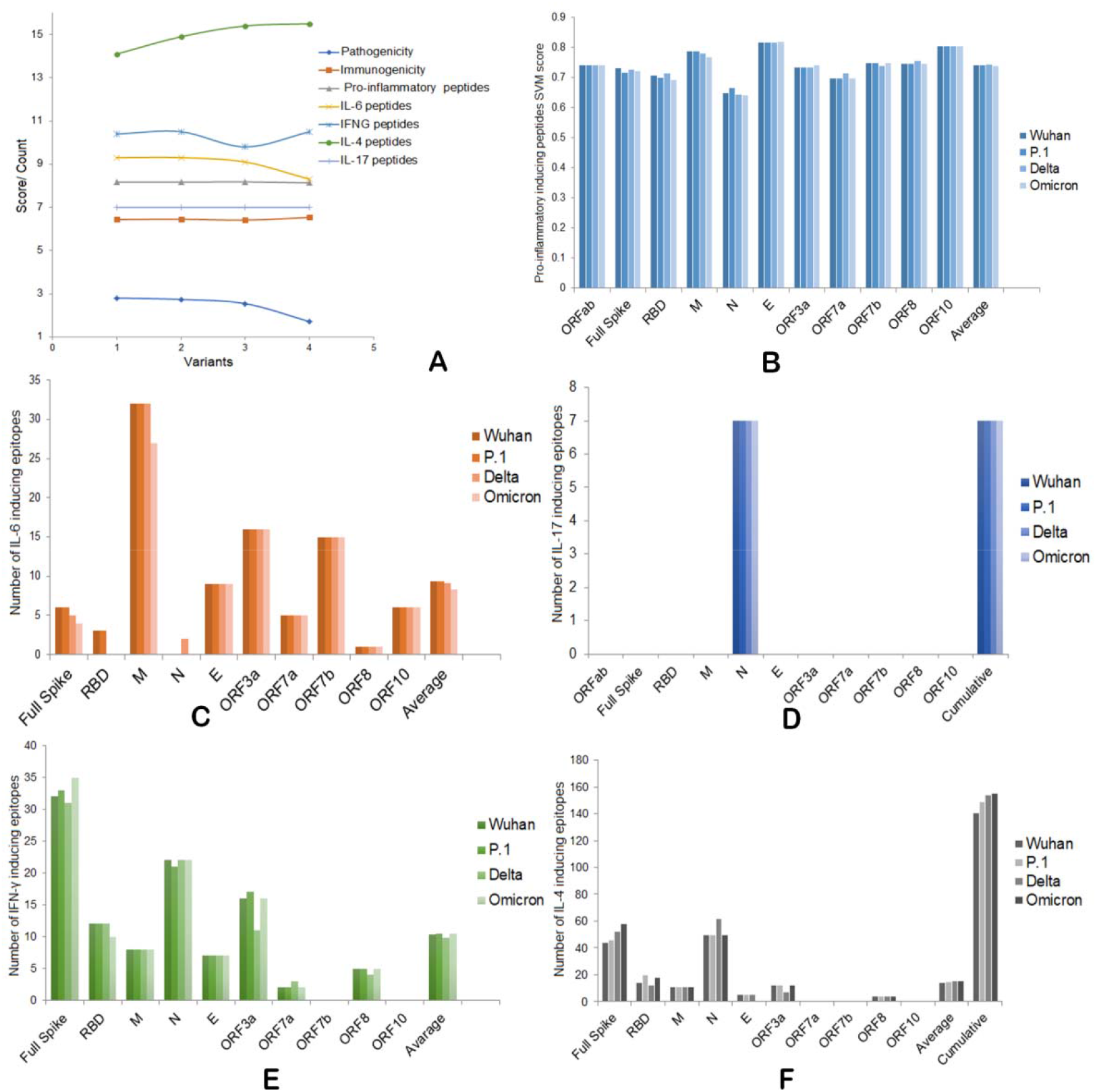
A) Overall pathogenic, immunogenic, IFNs, and ILs induction abilities of four SARS-CoV-2 variants. B) Pro-inflammatory epitope production scores of four variants. C) IL-6 inducing epitope counts of four variants. D) IL-17 inducing epitope counts of four variants. E) Number of IFN-γ inducing epitopes by four variants. F) Number of IL-4 inducing epitopes by four variants.

We found a varied antigenic potential of individual proteins of these variants, and there is an overall slight increase in the antigenic potential in the following order: Omicron > Gamma > Wuhan > Delta (**Fig 1D, 2A**). It is also important to note that the antigenic potential gradually increases in full-length S protein and spike RBD. Although a low antigenic score is observed for Omicron ORF3a compared to other variants, the antigenic scores of M, N, and ORF7a of Delta are low compared to other SARS-CoV-2 variants (**Fig 1D**). Taken together, our analysis suggests a decreased pathogenicity and increased immunogenicity for the Omicron variant.

### Omicron shows less pro-inflammatory and IL-6 producing epitopes as compared to other SARS-CoV-2 variants

Next, we focused on the efficacy of pro-inflammatory cytokine and IL-6 production ability by Omicron as compared to the other SARS-CoV-2 variants. In our analysis, at the individual protein level, we observed that RBD, ORF7a, and ORF8 of Delta showed increased pro-inflammatory inducing peptides scores compared to the other SARS-CoV-2 variants. The E protein had the highest score compared to other variants and the M protein showed a gradual decrease in the score in the order of Wuhan ≥ Gamma > Delta > Omicron (**Fig 2B**). Studying the overall pro-inflammatory inducing peptide score for the four variants, we observed the following order: Delta > Wuhan > Gamma > Omicron (**Fig 2A, B**). Therefore, the Omicron may produce less pro-inflammatory cytokines than the other variants.

We specifically focused on the IL-6 producing epitope counts and scores, and we observed the same trend in a very distinguished pattern. There is an overall gradual decrease in IL-6 inducing epitopes in the order: Wuhan ≥ Gamma > Delta > Omicron (**Fig 2A, C**). At individual protein level analysis, this trend was observed mainly for the full-length S protein and to some extent in the case of the M protein, where the Wuhan, Gamma, and Delta may produce an equal number (n=32) of IL-6 inducing epitopes; however, the Omicron M protein may produce a maximum of 27 epitopes (**Fig 2C**). While the full-length S protein produces IL-6-inducing epitopes for all variants, it is interesting to note that the spike RBD of the Delta and Omicron variants may not produce IL-6 inducing epitopes. On the other hand, the N protein of Delta may generate IL-6 inducing epitopes, but the N protein of other variants does not (**Fig 2C**). Therefore, as per our results, the N protein of Delta is important for its disease severity through induction of cytokine storm, and in Omicron, a low IL-6 induction may be associated with the reduced disease severity.

For IL-17 inducing epitope number and score, we did not see any differences across the variants. Only the N protein produces the total of seven IL-17 inducing epitopes with a score of 4.6 at cut-off > 0.5 for all variants; however their positions vary due to mutations (**Fig 2A, D**). Therefore, we presume that IL-17 may not have any specific role in disease susceptibility or severity in any of the SARS-CoV-2 variants we have used in this analysis.

### Omicron shows increased IFN-γ and IL-4 inducing epitopes than other variants

To understand these four variants’ anti-inflammatory cytokine production abilities, we analyzed the IFN-γ and IL-4 inducing epitopes by these four variants. The overall number and score of IFN-γ producing epitopes by Delta are less than the other variants, and there is very small difference in this number and score in the other three variants (**Fig 2A, E**). At the individual protein level, although the full-length Spike of Omicron generates more IFN-γ producing epitopes, its RBD shows less number of IFN-γ positive epitopes as compared to other variants. The ORF3a and ORF8 of Delta show less IFN-γ producing epitopes as compared to Omicron. However, the ORF7a of Delta produces more IFN-γ producing epitopes than other variants (**Fig 2E**). As per our analysis, the order of overall IFN-γ induction ability is Omicron = Gamma > Wuhan > Delta. These results suggest a possible role of low IFN-γ induction and disease severity by Delta variant.

For IL-4, we observed an opposite trend of IL-6. There is a gradual increase of overall IL-4 producing epitopes and scores in order: Omicron ≥ Delta > Gamma > Wuhan (**Fig 2A, F**). Spike, N, and ORF3a are the three main contributing proteins for this difference. While fulllength Spike of Omicron may produce 58 IL-4 inducing epitopes, this number is 44, 46, and 52 for Wuhan, Gamma, and Delta, respectively. While Wuhan, Gamma, and Omicron produce 50 epitopes, this number is 62 for Delta. For ORF3a, Delta may produce only seven IL-4 inducing epitopes, but the Wuhan, Gamma, and Omicron each produce twelve IL-4 inducing epitopes (**Fig 2A, F**). From these data, it is also evident that, IL-4 mediated anti-SARS-CoV-2 response is mainly regulated by Spike, N, and ORF3a and the mutations in these three proteins are involved in immune evasion by Delta variant and decreased severity in the case of the Omicron variant.

### SARS-CoV-2 M, N, and ORF8 protein variants may differentially regulate immune response by interacting with human STOM, G3BP1, DDX21, and GGH

Next, we attempted to understand which SARS-CoV-2 proteins regulate interferon and interleukin production through host protein interactions and the effects of mutation in these SARS-CoV-2 proteins in those interactions. From the literature, we found the N protein of SARS-CoV-2 induces IFN-γ expression [17,18], ORF8 regulates IL-17 signaling [20], and IFN-γ mediated antiviral responses [23], and M protein inhibits expression IFN-λ1 [24]. We modelled the mutant ORF8, M, and N proteins of the Omicron, Delta, and Gamma variants compared with the original Wuhan sequence.

For the M protein mutants, the LINCS L1000 ligand perturbations did not show any significant result (p>0.05) related to up or down-regulation of IFN or ILs through interaction with any human proteins. However, it is found that M protein may up-regulate IFN-γ, IFN-α, and IL-6 through interaction with human STOM (Stomatin), which regulate innate immunity. In protein-protein docking analysis, we found the binding scores for *h*STOM and M variants in this order: Delta ≥ Omicron > Wuhan = Gamma (**Fig 3A, B**). Therefore, the Delta and Omicron variants have more IFN-γ and IL-6 production than Wuhan and Gamma through *h*STOM mediated pathway. However, since the M protein and STOM mediated IFN-γ, IFN-α, and IL-6 production is not significantly (p>0.05) enriched in Enrichr analysis, we can ignore this finding (**Fig 3A, B**).

**Fig 3:**
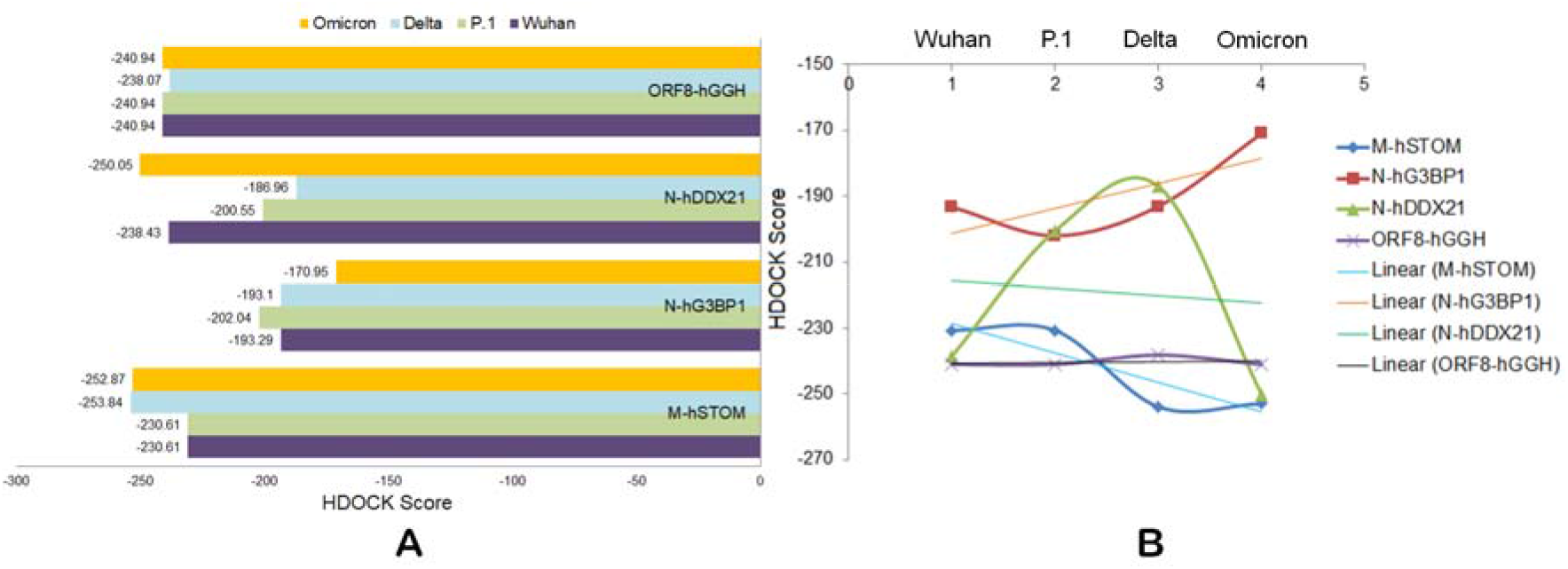
The binding strength between human STOM: M (SARS-CoV-2) variants, human G3BP1: N (SARS-CoV-2) variants, human DDX21: N (SARS-CoV-2) variants, and human GGH: ORF8 (SARS-CoV-2) variants. A-Bar chart and B-Scatter chart.

On the other hand, the N protein of SARS-CoV-2 was found to interact with human G3BP stress granule assembly factor 1 (G3BP1) and human DExD-Box helicase 21 (DDX21) with a significant p-value (p<0.05). These N: *h*G3BP1 and N: *h*DDX21 interactions are associated with increased anti-inflammatory IL-4 production in human cells. While we checked the binding of human G3BP1 with N protein variants, we observed the binding strength of the Omicron N protein is comparatively less than the N proteins from the other variants. The degree of the binding strength follows an order: Gamma > Wuhan ≥ Delta > Omicron. Therefore, we presume that the N protein of the Omicron variant may not increase the IL-4 production through *h*G3BP1 mediated pathway (**Fig 3A, B**). In contrast, the Omicron N protein was found to bind more strongly to the human DDX21 than the N proteins from other variants. Here the binding strength order is: Omicron > Wuhan > Gamma > Delta (**Fig 3A, B**). In Enrichr analysis, we found ORF8 of SARS-CoV-2 interacts with human GGH and down-regulates IL-17 and IFN-α. Further, the docking analysis suggests that mutations in ORF8 of the Delta variant slightly decrease the interaction between ORF8 and GGH. Therefore, the Delta variant may produce less IL-17 and IFN-α as compared to the other three variants (**Fig 3A, B**).

Taken together, our analysis suggests that strong interaction of the Omicron N protein with human DDX21 may be associated with increased anti-inflammatory IL-4 production in cases of Omicron infection that reduces the disease severity.

### Omicron Spike shows stronger binding to *h*ACE2, but its stability is low

To understand the increased transmutability of Omicron, we analyzed the binding strength between Spike RBD and *h*ACE2 as well as the stability of the various structural proteins of the four SARS-CoV-2 variants. The binding affinity (more negative score represents stronger binding) of the Spike-RBD to *h*ACE2 is highest for the Omicron variant compared to the other three SARS-CoV-2 variants. Both the HDOCK and Schrodinger-based PIPER pose scores show the same trend. PIPER pose scores-based order of binding affinity is: Omicron > Delta > Wuhan > Gamma, and the HDOCK-based order is: Omicron > Gamma > Wuhan > Delta (**Fig 4A, B**).

**Fig 4:**
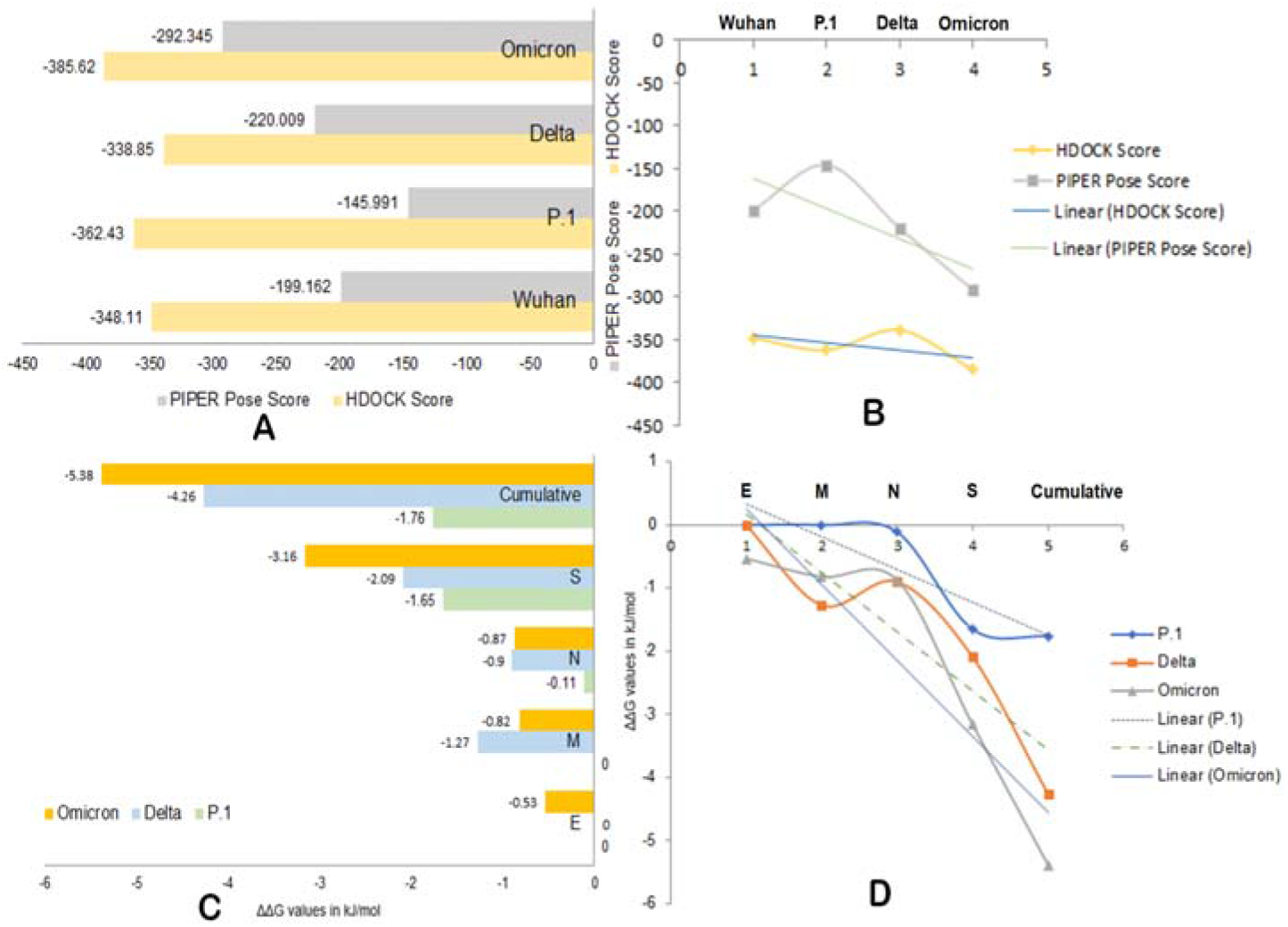
The binding strength of Spike-RBD and hACE2 of four SARS-CoV-2 variants (A-Bar chart and B-Scatter chart). The stability (ΔΔG values) analysis of SARS-CoV-2 structural proteins (A-Bar chart and B-Scatter chart).

To understand the stability of SARS-CoV-2 proteins from the four variants, we first identified the mutations in four key structural proteins (S, E, N, and M) of the Gamma, Delta, and Omicron variants compared with the amino acid sequence of the original Wuhan (wildtype) strain (RefSeq). The MSA-based identified mutations are presented in **Table-1**.

**Table 1:**
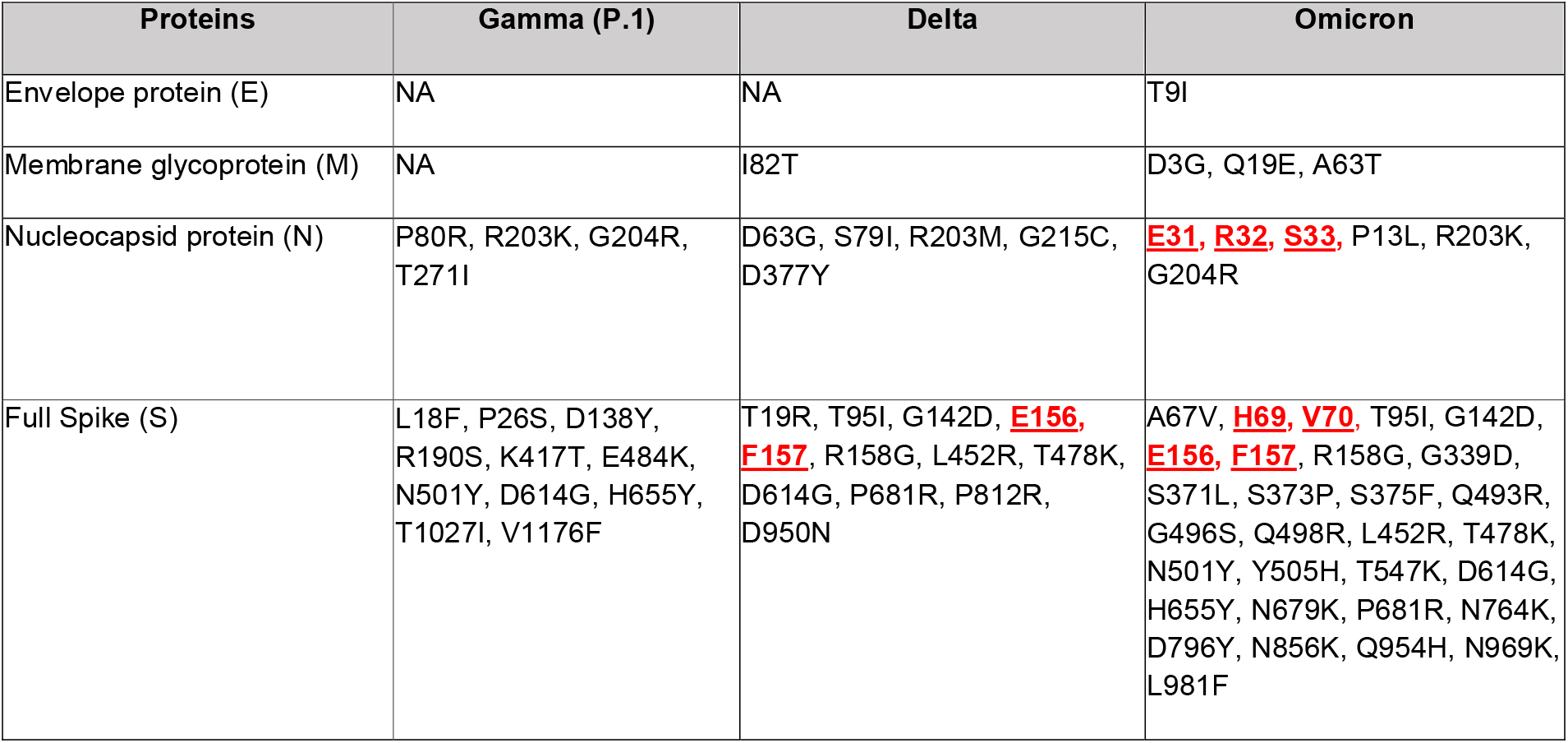
Mutations in structural proteins of Gamma, Delta, and Omicron compared to wildtype (Wuhan) strain of SARS-CoV-2. Bold red underlined residues are deleted residues.

| Proteins | Gamma (P.1) | Delta | Omicron |
| --- | --- | --- | --- |
| Envelope protein (E) | NA | NA | T9I |
| Membrane glycoprotein (M) | NA | I82T | D3G, Q19E, A63T |
| Nucleocapsid protein (N) | P80R, R203K, G204R, T271I | D63G, S79I, R203M, G215C, D377Y | **<u>E31</u>**, **<u>R32</u>**, **<u>S33</u>**, P13L, R203K, G204R |
| Full Spike (S) | L18F, P26S, D138Y, R190S, K417T, E484K, N501Y, D614G, H655Y, T1027I, V1176F | T19R, T95I, G142D, **<u>E156</u>**, **<u>F157</u>**, R158G, L452R, T478K, D614G, P681R, P812R, D950N | A67V, **<u>H69</u>**, **<u>V70</u>**, T95I, G142D, **<u>E156</u>**, **<u>F157</u>**, R158G, G339D, S371L, S373P, S375F, Q493R, G496S, Q498R, L452R, T478K, N501Y, Y505H, T547K, D614G, H655Y, N679K, P681R, N764K, D796Y, N856K, Q954H, N969K, L981F |

In mutation-based stability (ΔΔG values) analysis, all the four structural proteins of Omicron are unstable compared to the Gamma variant and the wild-type Wuhan strain. The M protein in the Omicron variant is more unstable than in the other three SARS-CoV-2 variants (**Fig 4C, D**). For the M protein, the Delta variant shows more instability compared to the Omicron variant. However, the stability of the M protein of the Gamma variant is the same as for the Wuhan strain (**Fig 4C, D**). The N protein of Omicron variant shows nearly the same degree of instability as the Delta variant. However, the instability of the N protein from the Gamma variant is very less as compared to the Omicron and Delta variants (**Fig 4C, D**). We observed a gradual decrease instability in the following order: Omicron > Delta > Gamma (**Fig 4C, D**). While we calculated the cumulative instability for all structural proteins, we observed that the Omicron proteins are more unstable, and the mutations in the S and E proteins mainly contribute to this increased instability (**Fig 4C, D**).

Taken together, our results suggest that the Omicron S protein binds to the *h*ACE2 more strongly as compared to other variants, and therefore, the transmission rate of the Omicron variant is higher. However, due to the higher instability of the S protein of the Omicron variant, the binding may not last long enough for systemic infection. Further, the E protein of the Omicron variant is most unstable and that may be associated with decreased viral particle production or maturation resulting in a lower viral load, severity, and faster recovery in case of Omicron infections.

## DISCUSSION

It is well established that the severity of disease and death rate from the Omicron variant are comparatively lower, but the transmissibility is higher than seen for any other SARS-CoV-2 variant [1–4]. According to our analysis, we found a gradual decrease in the pathogenic property of SARS-CoV-2 strains over time. The pathogenic property has decreased in the following order: Wuhan > Gamma > Delta > Omicron. Furthermore, in comparison to the Delta variant, the Omicron variant shows 20% and 32% lower pathogenicity at the genome and structural protein levels, respectively (**Fig 1A-C**). Recently, it was reported that the Omicron variant exhibits significant antigenic variation compared to other variants [50]. We also observed similar results in our sequence-based analysis. We found that the RBD, M, N, ORF3a, and ORF7a are the key proteins showing antigenic variation, whereas the Omicron RBD showed significantly higher antigenic properties than the other variants. The overall antigenic property found in our analysis is in the following order: Omicron > Gamma > Wuhan > Delta (**Fig 2A**). Our analysis also suggests that the Omicron variant could be the best possible attenuated vaccine, or its Spike RBD could be the best candidate to develop viral vector-, peptide-, and nucleic acid-based vaccines against COVID-19 in this current scenario.

It has been reported that the pro-inflammatory effect of the Omicron S protein is enhanced compared to other variants [51]. In our analysis, we found that the S protein may produce less pro-inflammatory and IL-6 inducing epitopes as compared to the Delta and other variants (**Fig 2B, C**). Further, the overall production of pro-inflammatory and IL-6 inducing epitopes are also lower in the Omicron variant than in other variants, and the order is Delta > Wuhan > Gamma > Omicron for pro-inflammatory epitopes and Wuhan ≥ Gamma > Delta > Omicron for IL-6 inducing epitopes. In addition to the S protein, the N protein plays an important role in production of these peptides (**Fig 2B, C**). On the other hand, we observed an increased anti-inflammatory IFN-γ and IL-4 induction ability of the Omicron variant compared to other variants in the following order: Omicron = Gamma > Wuhan > Delta and Omicron ≥ Delta > P.1Gamma > Wuhan, respectively (**Fig 2E, F**). We also noticed that the S, N, and ORF3a proteins play an important role in IFN-γ and IL-4 induction differences in these variants. Additionally we observed that the Omicron N protein could interact more strongly with human DDX21 than the equivalent protein of any other variant that may induce higher IL-4 production during Omicron infection (**Fig 3A, B**). The human DDX21 interacts with the SARS-CoV-2 N protein [38] and induces the innate immune response in dengue virus infection [52]. Taken together, these results suggest that the milder disease severity associated with Omicron infections may be related to its increased ability to induce IFN-γ and IL-4 and reduced ability to induce pro-inflammatory cytokines and IL-6 as compared to other variants.

Spike RBD binding to *h*ACE2 is the key event in SARS-CoV-2 infection and transmission, and it is believed that a strong RBD-*h*ACE2 interaction could be associated with hyper-transmissibility of the Omicron variant. However, conflicting reports have shown both strong [7–9] and weak binding affinity [10,11] of the Omicron RBD to *h*ACE2. Our analysis showed that the Omicron Spike-RBD had a stronger binding affinity to *h*ACE2 than the Spike-RBD of other SARS-CoV-2 variants, but that the Omicron S protein itself is more unstable than the corresponding S protein of other variants (**Fig 4C, D**). Although the strong affinity of the Omicron Spike-RBD to the *h*ACE2 may generate high transmissibility, the interaction may not be sufficient for systemic infection due to poor stability of Omicron S protein leading to severe COVID-19. Attenuated replication of Omicron is associated with decreased disease severity and reduced death rates, and the E protein plays an essential role in this replication attenuation [6]. We observed that the structural S, N, M, and E proteins of the Omicron variant are more unstable than the other variants, and importantly, the E protein of the Omicron variant is most unstable (**Fig 4C, D**). This unstable E protein may decrease the ability of replication or maturation of new viral Omicron, resulting in reduced viral load, disease severity, and faster recovery from Omicron infection.

## CONCLUSIONS

In this study, we analyzed four genome sequences, one from each of the Gamma, Delta, Omicron, and the Wuhan (wild-type) RefSeq. We analyzed four structural and six accessory proteins from these genome sequences using various bioinformatics approaches to understand why the Omicron variant is more transmissible but causes less severe disease. Our analyses revealed many critical biological mechanisms in these aspects and showed that the SARS-CoV-2 variants losing their pathogenicity and inflammatory cytokine production ability and increasing their immunogenic and anti-inflammatory cytokine induction ability. Using these mechanisms, over time, through mutations in major structural and non-structural proteins, SARS-CoV-2 is trying to adapt to an attenuated co-existence with its human host. If this trend continues, emerging SARS-CoV-2 variants may show enhanced transmissibility but causing an even milder COVID-19 than the Omicron variant today. The bioinformatics strategy used in this analysis may also be useful for understanding the dynamics of other viruses and may help being prepared for any other future pandemic caused by viral or bacterial infections.

## Author contribution

DB: Conceptualization, method development, data collection, analysis, and writing the paper, supervised and managed the project; ST: Performed mutation analysis; LGRG, CHRP: Performed epitope analysis; BSA, SA: Performed structural analysis; KJA, HJB, AAA: Performed reanalysis; AAA, SSH, MT, EMR, KR, VA: Provided technical inputs; KL, VNU: Provided technical inputs and edited the paper.

## Acknowledgment

VA and DB acknowledge the support from Coordenação de Aperfeiçoamento de Pessoal de Nível Superior (CAPES), Fundação de Amparo à Pesquisa do Estado de Minas Gerais (FAPEMIG), Conselho Nacional de Desenvolvimento Científico e Tecnológico (CNPq) and Pró-Reitoria de Pesquisa da Universidade Federal de Minas Gerais (PRPq). KJA acknowledges support from Taif University Researchers Program (Project Number: TURSP-2020/128), Taif University, Saudi Arabia.

